# Trait variation between and within Andes and coastal mountain ranges in the iconic South American tree *Araucaria araucana* in Chile

**DOI:** 10.1101/2022.01.04.474828

**Authors:** Mariah McIntosh, Jorge González-Campos, Patrick Demaree, Omayra Toro-Salamanca, Roberto Ipinza, Marcela A. Bustamante-Sánchez, Rodrigo Hasbún, Cara R. Nelson

## Abstract

As global commitments to restoration are underway, science is needed to support capacity to achieve meaningful gains for ecosystems and human communities. In Chile, identification and generation of appropriate plant material is a barrier to achieving major restoration goals under the Paris Climate Agreement. Understanding genetic differentiation among plant populations is needed to maximize restoration success. For *Araucaria araucana*, a highly threatened iconic South American tree, this information is greatly needed to guide restoration and conservation efforts because this species occurs across a strong climate gradient. We grew seedlings from 12 populations of *A. araucana* across its range in Chile in a common garden to assess regional (coastal versus Andes mountain ranges) and population variation in key plant traits and relate this variation to environmental variables. We demonstrate that *A. araucana* is differentiated within regions and populations across its range in Chile by a suite of traits, particularly branch number and length (showing plant architectural differences) and needle width (showing leaf investment differences). We show that this variation is at least partly explained by climate and soil variables, with the most variation explained by differences between regions in temperature annual range. Thus, we recommend that restoration efforts focus on conserving genetic variation among and within regions and their populations and preventing the translocations of genotypes between coastal and Andes populations.

## Introduction

As global ecosystems are increasingly affected by anthropogenetic degradation and climate change, ecological restoration is critically needed to repair ecosystems and support the human systems that depend on them. Towards that end, countries across the world are making ambitious restoration commitments. For instance, Chile aims to restore 1 million hectares of degraded land by 2050 as a part of its Nationally Determined Contribution under the Paris Climate Agreement (Gobierno de Chile 2020). One of the primary barriers to effective restoration is lack of understanding of appropriate plant materials (Gann et al. 2019; León-Lobos et al. 2020). To protect genetic diversity, avoid maladaptation to outplanting sites, and limit negative effects on adjacent populations, it is important to understand genetic differentiation among and within plant populations. (Lesica & Allendorf 1999; Kramer & Havens 2009; Breed et al. 2013). This information, however, is not yet available for many species of conservation concern in general, and specifically lacking in Chile, limiting restoration capacity (León-Lobos et al. 2020). We narrow this knowledge gap for the ancient and iconic South American conifer, *Araucaria araucana* (pewen), a tree of high cultural and ecological value in South America. Most genetic information for this threatened species addresses neutral genetic variation (e.g., Souza et al. 2008, Martín et al. 2014), thus we lack information on adaptive genetic variation (Bekessy et al. 2003). Here, we characterized among- and within-population variation in key plant traits across the range of pewen in Chile and related overall trait variation to climate and soil variables, which commonly drive large-scale patterns of differentiation in trees (Alberto et al. 2013). Our work provides the basis for both understanding patterns of genetic and phenotypic variation across the range of this species and improving management and restoration capacity.

As plants are rooted in place and cannot escape environments in which they germinate, they are often adapted to local conditions and thus genetically and phenotypically differentiated by environment across their ranges (Leimu & Fischer 2008; Anderson et al. 2011). As a result, population differentiation is extremely common in plants (Leimu & Fischer 2008) and occurs across spatial scales from meters (Lekberg et al. 2012) to hundreds of kilometers (Liepe et al. 2016; Supple et al. 2018). For instance, population differentiation has been found in 90% of forest trees studied (Alberto et al. 2013). It is not surprising that local adaptation is so common, as it has been shown to improve plant growth, reproduction, and survival at home sites (Joshi et al. 2001; Leimu & Fischer 2008). If plants are moved to foreign environments outside their range of local adaptation, population fitness may be low and deleterious effects may occur in adjacent populations (Lesica & Allendorf 1999; Hufford & Mazer 2003; McKay et al. 2005; Broadhurst et al. 2008). Thus, understanding genetic differentiation among and within populations of the same species in key fitness traits is critical to informing conservation and restoration across the species range (Hufford & Mazer 2003; Broadhurst et al. 2008; Breed et al. 2013; Gann et al. 2019).

Beyond characterizing patterns of population differentiation, there is considerable interest in identifying environmental variables that explain these patterns (Reich et al. 1997; Wright et al. 2005; Alberto et al. 2013; Aitken & Whitlock 2013; Anderegg et al. 2016, 2018). Climate gradients are often considered as drivers of plant population differences (Alberto et al. 2013; Bower et al. 2014), as plant distribution is strongly driven by climate (Webb 1986; Woodward 1987; Woodward et al. 2004). As a result of provenance studies which have been conducted for multiple centuries, within species, climate variably explains population differentiation depending on species, traits studied, and the magnitude of climate gradients (Alberto et al. 2013; Griffin-Nolan et al. 2018). Soil variables may play a role in driving population differentiation that is equal to or even greater than that of climate, despite soil variables varying at much smaller spatial scales (Macel et al. 2007; Lekberg et al. 2012; Siefert et al. 2014; Lajoie & Vellend 2015; Gibson et al. 2019). However, the relative contribution of these factors (and the scale of their variation) remains unresolved (but see Siefert et al. 2015). Here, we ask which climate and soil variables best explain multivariate genetic trait differentiation among populations, addressing large-scale climate versus small-scale soil heterogeneity as drivers of population differentiation.

There is increasing recognition of the importance of maintaining both genetic and phenotypic variation in species-specific conservation and restoration strategies, especially given anticipated rapid changes in climate (Kramer & Havens 2009; Breed et al. 2013; Havens et al. 2015; Gann et al. 2019). Understanding this genetic variation is valuable for managers as genetic variation can be both the result of previous natural selection and the raw material for future selection in response to environmental change (Kramer & Havens 2009; Kremer et al. 2012). Furthermore, understanding the extent to which within-species variation occurs within or among populations (population versus regional variation) may have implications for the appropriate sourcing of genetic material for restoration. For example, in a study of the threatened species *Eucalyptus melliodora* in Australia, most genetic variation occurred within versus among populations, and the authors concluded that seeds could be sourced broadly for restoration (Supple et al. 2018). Similarly, a high level of within-population variation was identified for a relatively small number of locally adapted populations of interior spruce complex *(Picea glauca, P. engelmannii*, and their hybrids) and lodgepole pine (*Pinus contorta)* across an area spanning British Columbia and Alberta (>1000 km in latitude and longitude) in Canada (Liepe et al. 2016). Meta-analysis supports these case studies to show that for trees (particularly those that are wind pollinated), this pattern of population differentiation across large spatial scales (on the order of hundreds to thousands of kilometers) and high within-population variation is common, even when gene flow is significant (Savolainen et al. 2007; Alberto et al. 2013; Liepe et al. 2016). However, the majority of this information is for temperate forest trees with large ranges (Alberto et al. 2013) and we don’t yet know how species with restricted and fragmented ranges vary among and within populations.

Although understanding drivers and spatial patterns of genetic and phenotypic variation is generally important for ecosystem management, it is particularly important to have this information for pewen. There is considerable interest in restoration of this species across its range and restoration programs are in progress, but lack of information on genetically-based phenotypic variation (rather than neutral genetic variation, which has been largely resolved; see (Martín et al. 2014) limits understanding of genetically appropriate material for outplanting and ability to conserve genetic diversity (León-Lobos et al. 2020). Additionally, this species is experiencing drought-related mortality that varies among and within regions (Willhite 2019; Puchi et al. 2021), suggesting that climate and soil conditions may predict survival outcomes and adding urgency to the need for information on regional and population differentiation for this species.

We studied patterns of among- and within-population genetic variation of pewen across its range in Chile, in order to improve both ecological understanding and management and restoration of this unique species. Our study is one of only a handful that addresses within-species genetic variation in a suite of traits rangewide in South American conifers. For pewen, we build on previous phenotypic and genetic studies in this species that were limited in the number of sites and traits sampled to assess among- and within-population variation in a broad suite of traits and relate this variation to climate and soil variables. Specifically, we assessed: whether plants from populations that experience different climate and soil conditions show trait variation among or within populations and regions (Andes vs. coastal mountain ranges) (Q1); which plant traits drive overall differences in phenotypes among and within populations and regions (Q2); and which climate and soil variables drive overall differences in phenotypes among and within populations and regions (Q3). Our findings contribute to the growing literature on among- and within-population variation in trees and uses common methods for developing seed transfer guidelines to lay the groundwork for developing these important resources for this species.

## Methods

### Study System

*Araucaria araucana* (pewen) is native to the coastal and Andean cordilleras of central Chile (37° 31’ to 39° 30’) and Argentina (37 ° 45’ to 40° 20’) (Aagesen 1998; Figure 1). The range of pewen, although relatively small, spans substantial elevation (664-1227 m), precipitation (1100-2219 mm annual precipitation), and temperature (6.1-9.6 °C mean annual temperature) gradients (Table 1). *A. araucana* is a dioecious and wind pollinated masting species (Sanguinetti and Kitzberger 2008). This species is of cultural and spiritual importance to the Mapuche Pewenche (pewen people), and the sale and consumption of *ngülliw* (the large pinenut-like seeds of pewen) is important for subsistence (Herrmann 2006). Pewen has been listed as “Endangered” on the IUCN Red List since 2011 (Premoli 2015) due to historic deforestation (although it is now protected by the government of Chile), invasion by *Pinus contorta* (lodgepole pine), illegal harvest of seeds (legal for indigenous peoples only), and seed consumption by livestock (Cóbar-Carranza et al. 2014; Premoli 2015; Tella et al. 2016). Seed regeneration is poor, but vegetative reproduction may occur (Aagesen 1998). Because its significant climate gradient in Chile and its ecological and cultural importance, pewen is an excellent study system for addressing management-relevant questions about patterns and predictors of genetic variation among and within populations across a species’ range.

**Figure 1.**
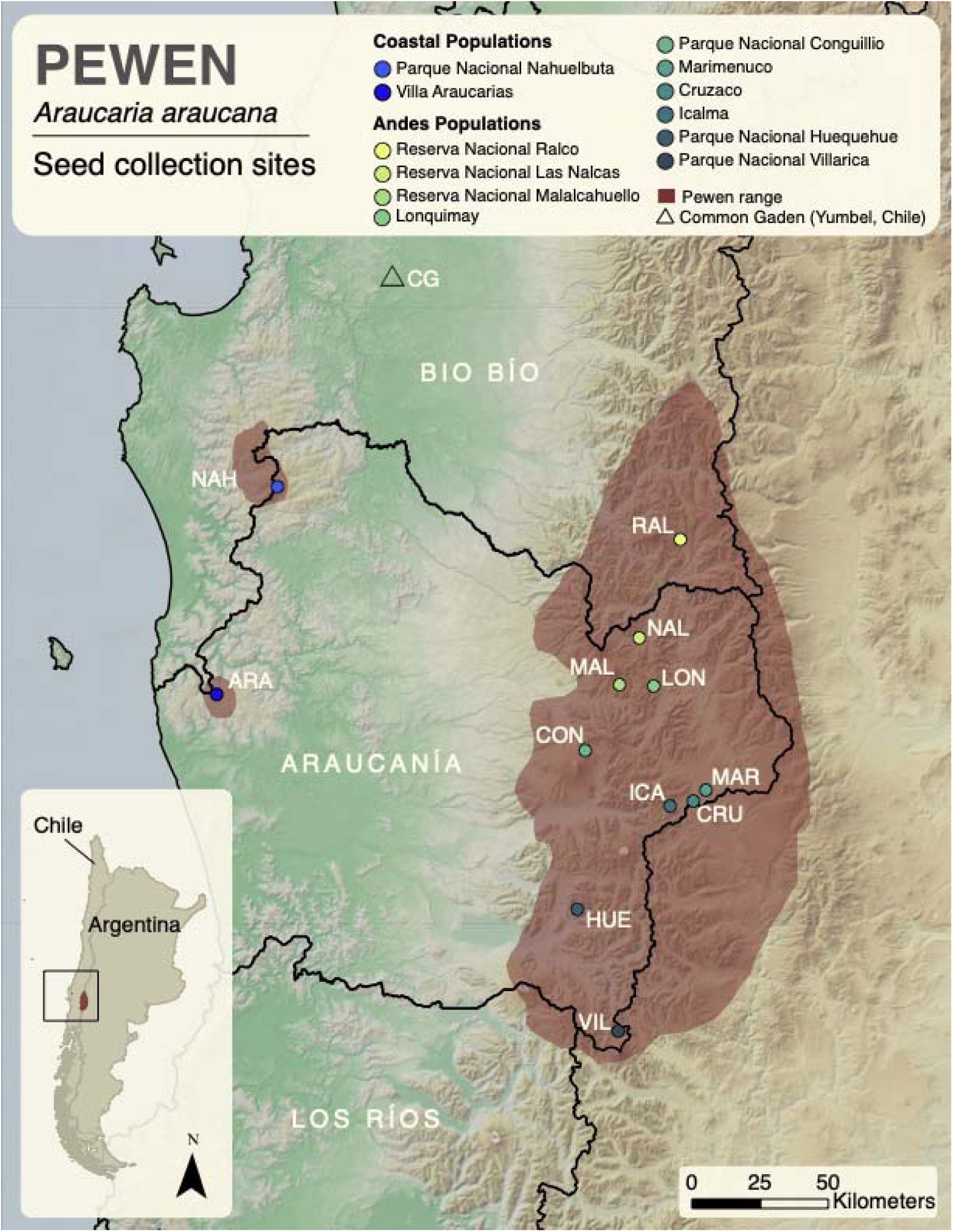
Map of seed collection sites in Chile. The range of pewen is shown in brown. Study populations are shown by dots, with colors corresponding to populations as used in subsequent figures. The common garden site (Yumbel, Chile) is labeled with a triangle.

**Table 1.** Collection sites covering the range of pewen in Chile vary in altitude and climate variables.

| **Population** | **Region<sup>1</sup>** | **Code<sup>2</sup>** | **Families<sup>3</sup>** | **Latitude** | **Longitude** | **Alt.<sup>4</sup>** | **MAP<sup>56</sup>** | **MAT<sup>7</sup>** | **TAR<sup>8</sup>** |
| --- | --- | --- | --- | --- | --- | --- | --- | --- | --- |
| *PN<sup>9</sup> Nahuelbuta* | Coastal | NAH | 20 | -<br>37.805609 | -<br>73.017985 | 1269 | 1604 | 6.1 | 19.9 |
| *Villas Araucarias* | Coastal | ARA | 20 | -<br>38.495328 | -<br>73.254247 | 664 | 1501 | 8.9 | 19.4 |
| *RN<sup>10</sup> Ralco* | Andes | RAL | 20 | -<br>37.939491 | -<br>71.334323 | 1239 | 1696 | 8.8 | 25.4 |
| *RN Las Nalcas* | Andes | NAL | 20 | -<br>38.269358 | -<br>71.489768 | 976 | 2219 | 9.6 | 24.7 |
| *RN Malalcahuello* | Andes | MAL | 20 | -<br>38.425844 | -71.56517 | 1382 | 1765 | 7.7 | 24.3 |
| *Lonquimay (\*)* | Andes | LON | 7 | -<br>38.426427 | -<br>71.421637 | 1376 | 1632 | 8.1 | 24.7 |
| *PN Conguillio* | Andes | CON | 19 | -<br>38.647372 | -<br>71.698783 | 1236 | 1860 | 7.9 | 23.6 |
| *PN Huequehue* | Andes | HUE | 20 | -<br>39.172097 | -<br>71.707628 | 1378 | 1464 | 6.7 | 23.0 |
| *Cruzaco* | Andes | CRU | 20 | -<br>38.800124 | -<br>71.235559 | 1424 | 1138 | 7.9 | 24.7 |
| *Icalma* | Andes | ICA | 20 | -<br>38.819732 | -<br>71.332654 | 1195 | 1355 | 8.7 | 24.4 |
| *Marimenuco (\*)* | Andes | MAR | 7 | -<br>38.762456 | -<br>71.184824 | 1401 | 1110 | 8.2 | 24.7 |
| *PN Villarica* | Andes | VIL | 20 | -39.569 | -<br>71.514951 | 1187 | 1145 | 7.5 | 23.0 |

<sup>1</sup> Region refers to the mountain range from which the populations were selected (Andes vs. Coastal range, see Figure 1).
<sup>2</sup> Codes refers to populations throughout the manuscript.
<sup>3</sup> Families refer to sample size for each population.
<sup>4</sup> Alt. = altitude (m)
<sup>5</sup> Climate variable means were calculated by accessing WorldClim data for each family's latitude and longitude coordinates (tree from which seeds were sampled) and calculating means at the population level.
<sup>6</sup> MAP = mean annual precipitation (mm)
<sup>7</sup> MAT = mean annual temperature (°C)
<sup>8</sup> TAR = temperature annual range (maximum temperature – minimum temperature; °C)
<sup>9</sup> PN = Parque Nacional
<sup>10</sup> RN = Reserva Nacional

### Study Populations

We selected 12 sites (referred to hereafter as populations) throughout the range of pewen in both the Andean and coastal mountain ranges (regions) of Chile spanning altitude and climate gradients (Table 1, Figure 1). Populations were located within five genetic clusters (two coastal, three Andean) identified by Martín et al. (2014) using a landscape genetic approach.

### Seed Collection, Seedling Growth, and Trait Measurements

At each population, we collected seeds from trees that were at least 150 m apart and had available seeds in 2018 at the time of collection. Trees for seed collection were not chosen randomly, as they had to be producing seeds, and many were chosen nearby roads or trails because of convenient access (see *Limitations* in Discussion). We referred to seeds from a single tree at a given population as a family (specifically, they are half sibling families). At each population, we initially sampled 7-96 families per population depending on site size and availability; we randomly selected 20 families from each of the 12 populations for inclusion in the study (n=1 seedling per family), except in two populations (Lonquimay and Marimenuco), where n=7. Additionally, one individual was not measured by accident, reducing n to 19 for this population (Table 1).

After cold stratification at 4°C for two months, we cut the end of each seed and submerged them in water for 2 days at 4°C. Seeds were planted in plastic flats, germinated in a greenhouse in Yumbel, Chile (−37.098090, -72.562230), and then grown for one year. Seedlings experienced ambient light conditions and were well-watered (at least once and sometimes more than twice per day depending on temperature). We were unable to randomize the location of individual seedlings on benches because of the requirements of the commercial growing facility; however, we anecdotally noted that the effects of population and family on seedling traits were more prominent than greenhouse effects (see *Limitations* in Discussion). Germination rate was measured at 30 weeks, and seedling survival was measured after one year. Plants that were fully browned were considered dead.

In December of 2019, we measured a suite of traits to assess variation among and within regions and populations. Because no information exists on which traits are adaptive for this species, we selected traits related to seedling growth and biomass allocation, architecture, and leaf economics that are known to relate to resource use and stress-tolerance strategies (Table 2). We counted the number of whorls (opposite branches originating from a single point) and branches of each seedling and measured stem length, basal diameter, and the length of each branch (to calculate a mean branch length; if there were no branches, branch length was 0). Additionally, we measured the length and width of the three longest needles to calculate maximum needle lengths and widths (referred to as needle length and width throughout). We measured needle area using the app LeafByte (Getman-Pickering et al. 2020) and calculated needle mass per area. To measure needle density, we measured needle volume using the water displacement method (Hughes 2005) and divided volume by needle mass. Needle thickness was calculated by dividing needle volume by needle area. Additional descriptions of trait measurements and units are included in Table 2.

**Table 2.**
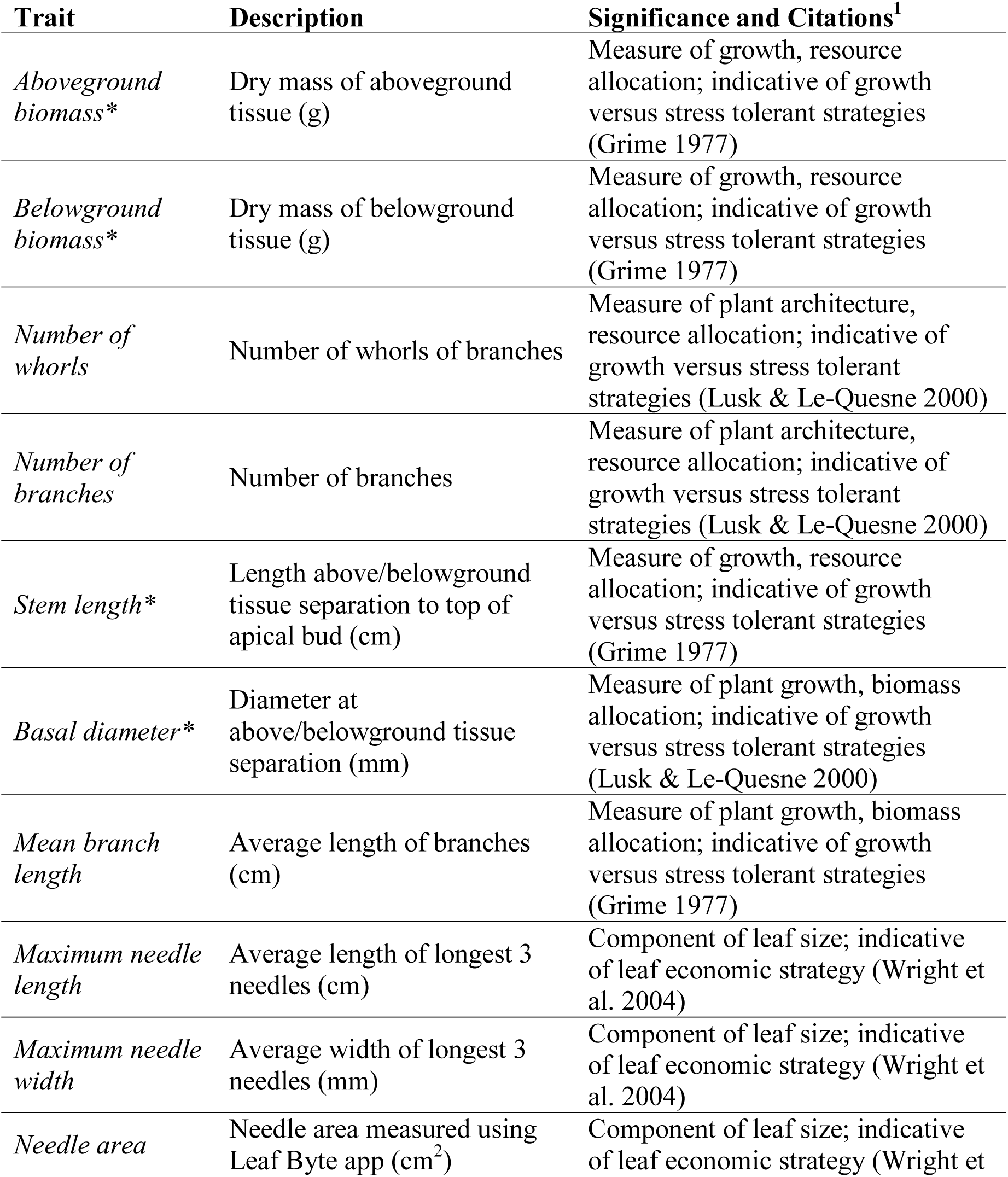
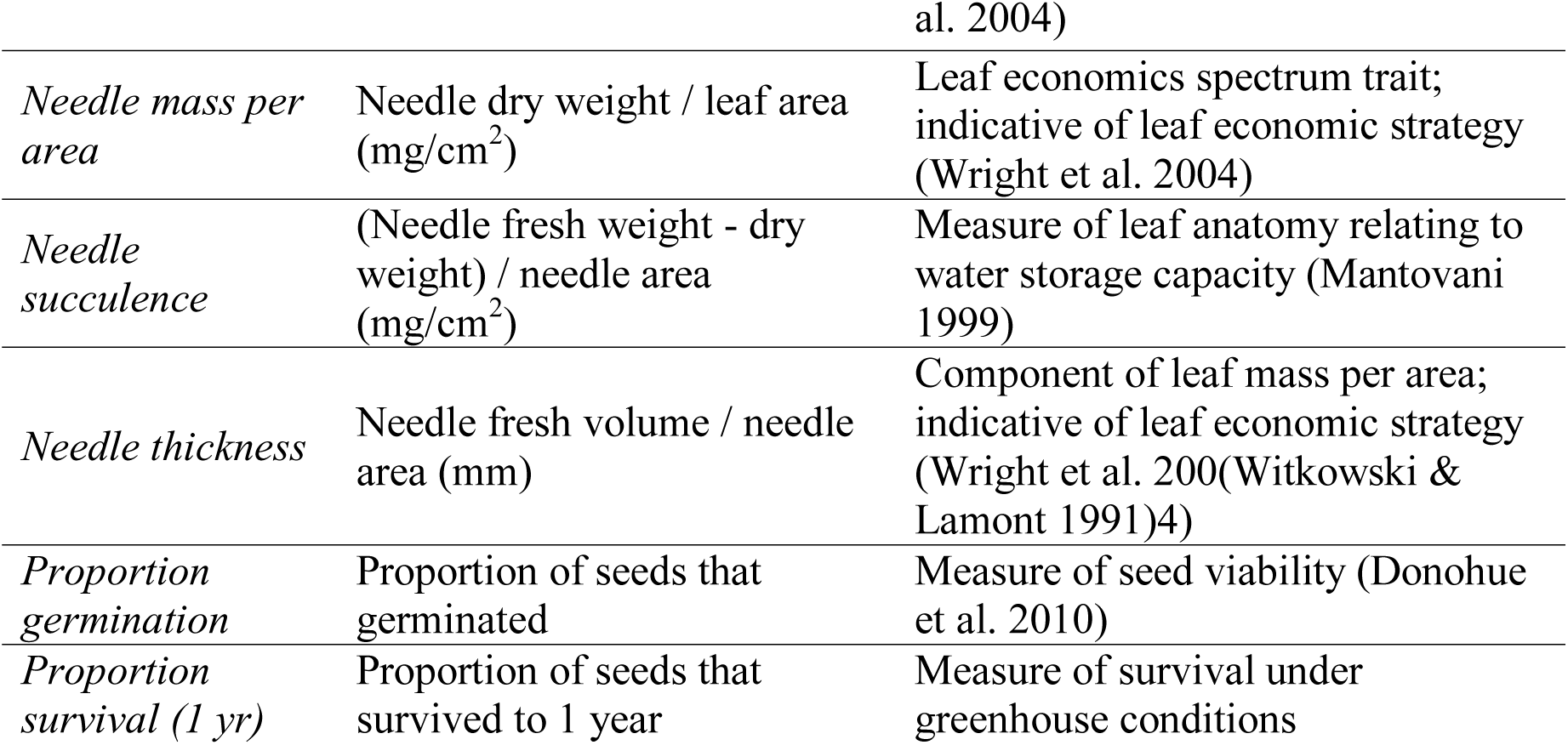
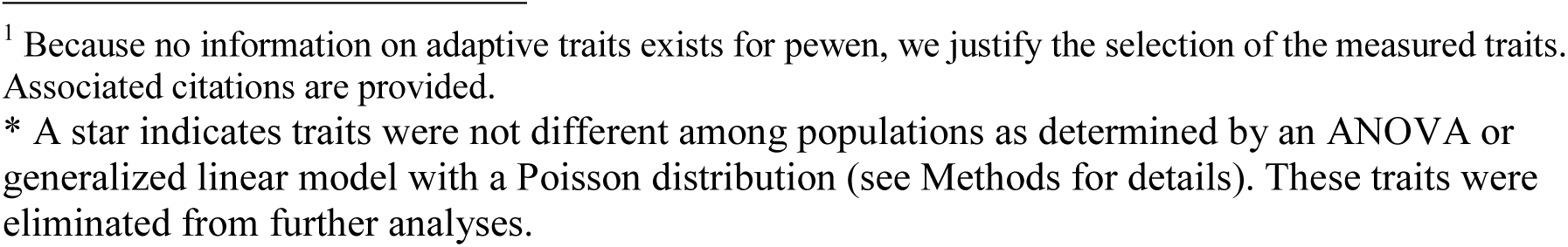
Traits measured in common garden seedlings.

| **Trait** | **Description** | **Significance and Citations<sup>1</sup>** |
| --- | --- | --- |
| *Aboveground biomass\** | Dry mass of aboveground tissue (g) | Measure of growth, resource allocation; indicative of growth versus stress tolerant strategies (Grime 1977) |
| *Belowground biomass\** | Dry mass of belowground tissue (g) | Measure of growth, resource allocation; indicative of growth versus stress tolerant strategies (Grime 1977) |
| *Number of whorls* | Number of whorls of branches | Measure of plant architecture, resource allocation; indicative of growth versus stress tolerant strategies (Lusk & Le-Quesne 2000) |
| *Number of branches* | Number of branches | Measure of plant architecture, resource allocation; indicative of growth versus stress tolerant strategies (Lusk & Le-Quesne 2000) |
| *Stem length\** | Length above/belowground tissue separation to top of apical bud (cm) | Measure of growth, resource allocation; indicative of growth versus stress tolerant strategies (Grime 1977) |
| *Basal diameter\** | Diameter at above/belowground tissue separation (mm) | Measure of plant growth, biomass allocation; indicative of growth versus stress tolerant strategies (Lusk & Le-Quesne 2000) |
| *Mean branch length* | Average length of branches (cm) | Measure of plant growth, biomass allocation; indicative of growth versus stress tolerant strategies (Grime 1977) |
| *Maximum needle length* | Average length of longest 3 needles (cm) | Component of leaf size; indicative of leaf economic strategy (Wright et al. 2004) |
| *Maximum needle width* | Average width of longest 3 needles (mm) | Component of leaf size; indicative of leaf economic strategy (Wright et al. 2004) |
| *Needle area* | Needle area measured using Leaf Byte app (cm<sup>2</sup>) | Component of leaf size; indicative of leaf economic strategy (Wright et |

<sup>1</sup> Because no information on adaptive traits exists for pewen, we justify the selection of the measured traits. Associated citations are provided.
\* A star indicates traits were not different among populations as determined by an ANOVA or generalized linear model with a Poisson distribution (see Methods for details). These traits were eliminated from further analyses.

|  |  |  |
| --- | --- | --- |
| *Needle mass per area* | Needle dry weight / leaf area (mg/cm<sup>2</sup>) | Leaf economics spectrum trait; indicative of leaf economic strategy (Wright et al. 2004) |
| *Needle succulence* | (Needle fresh weight - dry weight) / needle area (mg/cm<sup>2</sup>) | Measure of leaf anatomy relating to water storage capacity (Mantovani 1999) |
| *Needle thickness* | Needle fresh volume / needle area (mm) | Component of leaf mass per area; indicative of leaf economic strategy (Wright et al. 200(Witkowski & Lamont 1991)4) |
| *Proportion germination* | Proportion of seeds that germinated | Measure of seed viability (Donohue et al. 2010) |
| *Proportion survival (1 yr)* | Proportion of seeds that survived to 1 year | Measure of survival under greenhouse conditions |

### Climate and Soil Variables

We accessed climate and soil variables from WorldClim (Fick & Hijmans 2017), TerraClimate (Abatzoglou et al. 2018), and SoilGrids (Hengl et al. 2017) databases for the GPS coordinates of each family (see Table 3 for variables and units). WorldClim data were downloaded directly into R using the getData() function in the package raster (Hijmans & Van Etten 2021) in R Studio version 1.2.5042 (RStudio Team 2020). We extracted data for our coordinates using the extract() function in the package sp (Pebesma & Bivand 2005). For TerraClimate data, we used the getTerraClim() function in climateR (Johnson 2020) to download and extract data for our populations. Additionally, we accessed climate variables using regional climate models from the Center for Climate and Resilience Research at the Universidad de Chile (CR2; http://www.cr2.cl/) but they did not perform better than data from global models, so we excluded them from final analyses. For SoilGrids data, we used Google Earth Engine to access variables listed in Table 3.

**Table 3.**
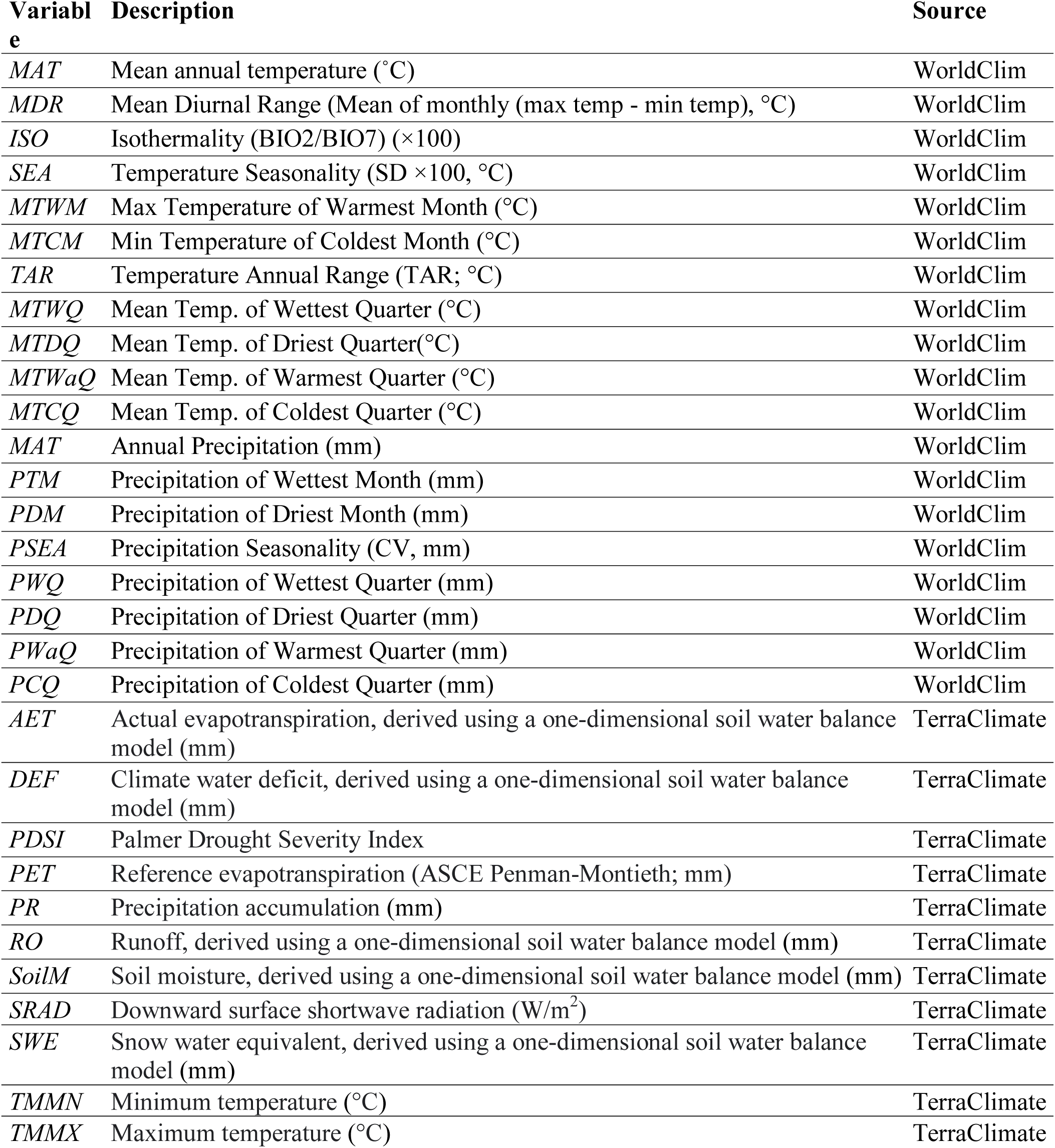
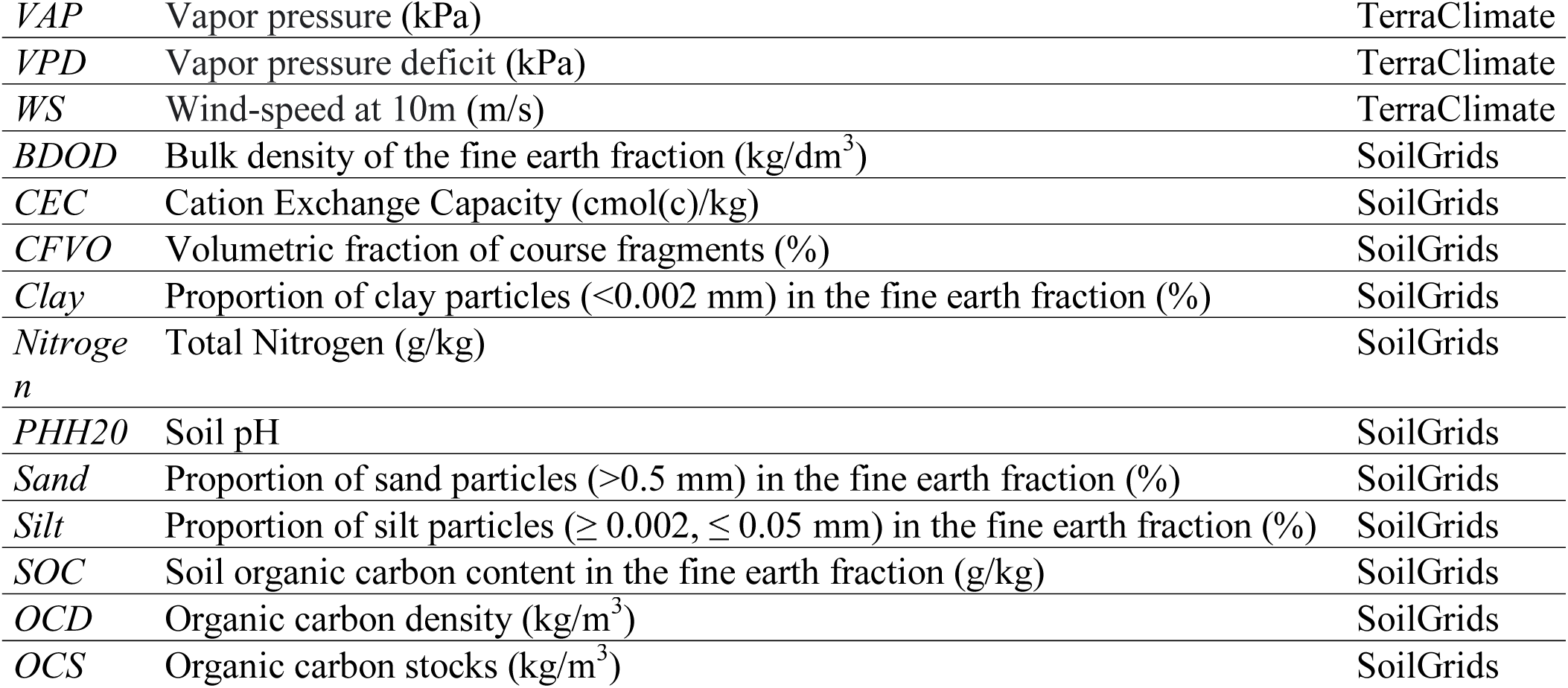
Bioclimatic and soil variables used for multiple regressions with trait PC scores (units in parentheses) extracted for the latitude and longitude coordinates of each tree from which seeds were sampled.

### Statistical Analysis

As our traits were measured in a common garden, which controls for most environmental variation, we assume trait differences are due to genetic differences rather than environment. To assess if individual traits varied among regions (coast vs. Andes) and populations (Q1), we used analysis of variance (ANOVA). For each trait, we ran nested models with region and population nested within region as factors to address the relative contribution of region and population and to identify traits which varied among populations and should be included in additional analysis (Supporting Information). Assumptions of ANOVA were checked using residuals plots and normal quantile plots. For count traits only (number of whorls, number of branches), we used a generalized linear model with a Poisson distribution instead of an ANOVA because these traits were not normally distributed (O’Hara & Kotze 2010). Traits that did not significantly vary among populations or regions (p>0.05) were not used for additional analyses (see Table 2 and Supporting Information for a list of the eliminated traits).

To address multivariate trait differences among regions and populations (Q1), we used principal component analysis (PCA) using Bray-Curtis distances with pairwise deletion of missing observations in the vegan package in R (Oksanen et al. 2019). We used multiple regressions with PC scores as response variables and traits as predictors to address which traits best explained overall differences in phenotypes regions and populations (Q2). A separate model was created for each of the first four PC axes (which explained 94% of variation). To select traits to include in our models, we used Spearman’s r to identify the traits most correlated with each axis where |r| ≥ 0.20 and p ≥ 0.05 (Supporting Information). We then excluded traits that covaried with other traits using the cutoff of |r|>0.60 (Zuur et al. 2010), selecting traits with higher correlation with axes scores first and removing less highly correlated traits that covaried. We used backwards selection to remove additional traits that did not add predictive power to the model using the step() function in R (RStudio Team 2020).

To address which climate and soil variables best explained overall differences in phenotypes among regions and populations (Q3), we created separate multiple regression models for each PC axis using climate and soil variables as predictors. For each of the first three PC axes, we used Spearman’s r to identify the traits most correlated with each axis where |r| ≥ 0.20 and p ≥ 0.05 (Supporting Information). We used backwards selection to remove additional traits that did not add predictive power to the model (p > 0.05). Overall contribution of climate variables in explaining trait variation across axes was assessed using PERMANOVA with the adonis() function in vegan with pairwise deletion of missing observations (Oksanen et al. 2019).

All analyses were conducted using RStudio version 1.2.5042 (RStudio Team 2020), and all figures except Figure 1 were made in R using ggplot2 (Wickham 2016). Figure 1 was made in ArcMap.

## Results

### Pewen seedlings from across regions and populations range-wide differ in their traits (Q1)

Plants from different regions (coast and Andes) and populations varied significantly in their traits (Figure 2, Supporting Information). Across the 16 measured traits, 11 differed significantly to varying degrees among regions and populations (Figure 2A, Supporting Information). Across all traits, regions were highly distinct, with coastal populations differing from Andes populations (Figure 2B) both in PC1 (79.3% of overall variation) and PC2 (9.6% of overall variation). Populations within each region also varied significantly in their traits. For PC1, region accounted for 9.8% of variation in axis scores (p<0.001) and population accounted for 12.1% of variation (p<0.001) in a nested ANOVA model. For PC2, region accounted for 8.9% of variation (p<0.001), and population accounted for 1.9% of variation (p<0.001).

**Figure 2.**
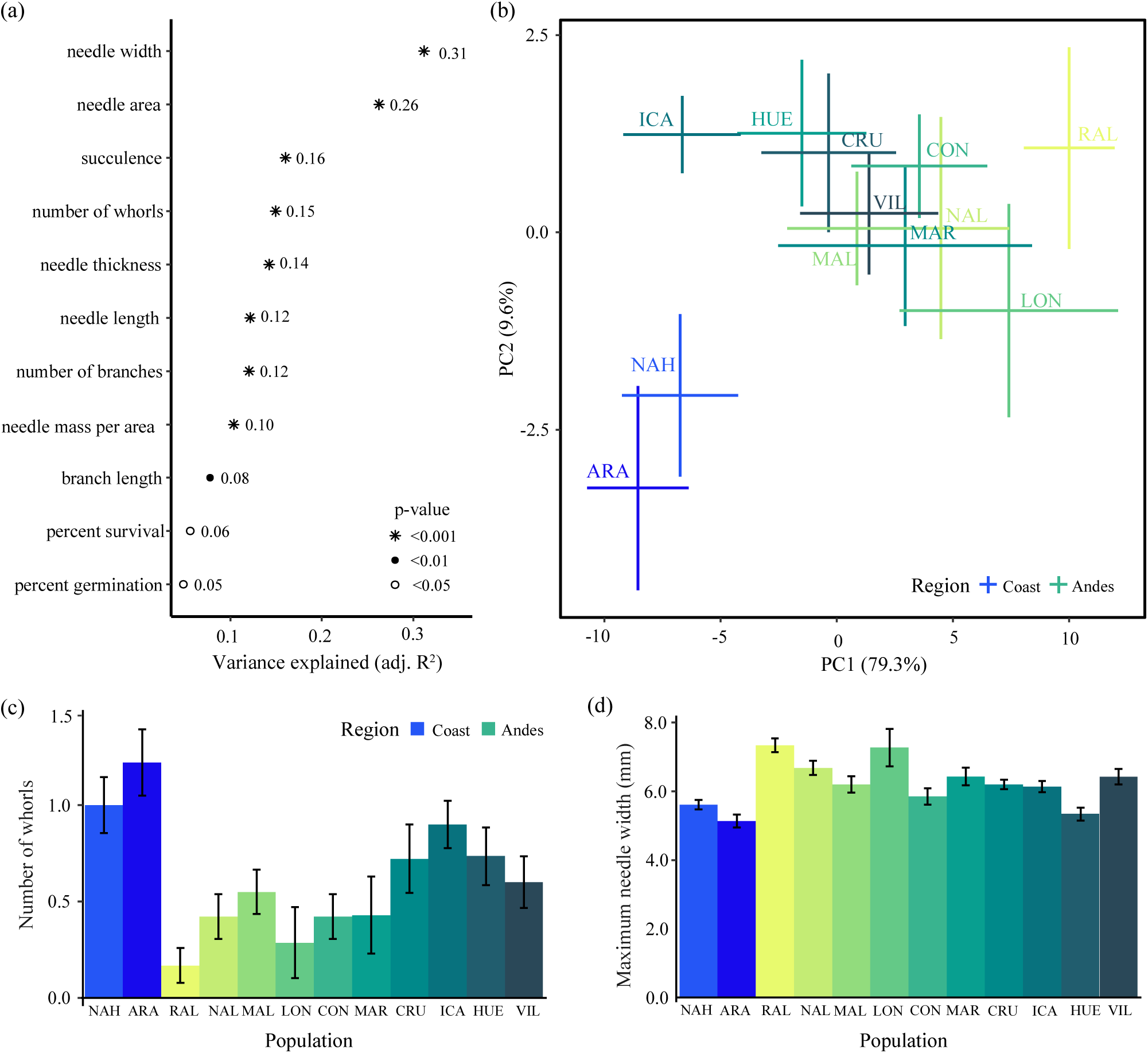
**A)** A suite of traits varies significantly among regions and populations across the range of pewen in Chile. Adjusted R^2^ values (numbers) and p-values (symbols, see legend) for ANOVA models of plant traits by populations. Only traits for which p<0.05 are included (see Table 2 for excluded traits). See Table 2 for trait units. **B)** Traits of pewen vary by region and population. Mean PCA scores for PCA axes 1 and 2 (percent variation explained in parentheses) for each population (3-letter codes, see Table 1, Figure 1). Bars show standard errors. **C)** Branch and needle traits vary among and within regions and populations across the range of pewen in Chile. Bars show standard errors. Color categories correspond to region (green = Andes, blue = coast), with gradients by latitude from north (light) to south (dark). Population codes are printed (see Table 1 for more information on populations).

### Branch architecture and needle traits explain overall region and population trait differences (Q2)

Branch architectural and needle traits explain overall trait differences between regions and among populations. For PC1, number of whorls, needle area, and needle succulence explained overall trait variation (Adjusted R^2^ = 0.83, p<0.001; Table 4, Supporting Information). Our model initially included proportion of survival to 1 year; but it did not provide explanatory power beyond included variables (and was removed per our backwards selection method; ΔAIC=1.7; Supporting Information). The number of branches and branch length were both highly correlated with the number of whorls (and thus not included in the model; Supplemental Information); and showed similar patterns among populations as number of whorls (shown in Figure 2C).

**Table 4.**
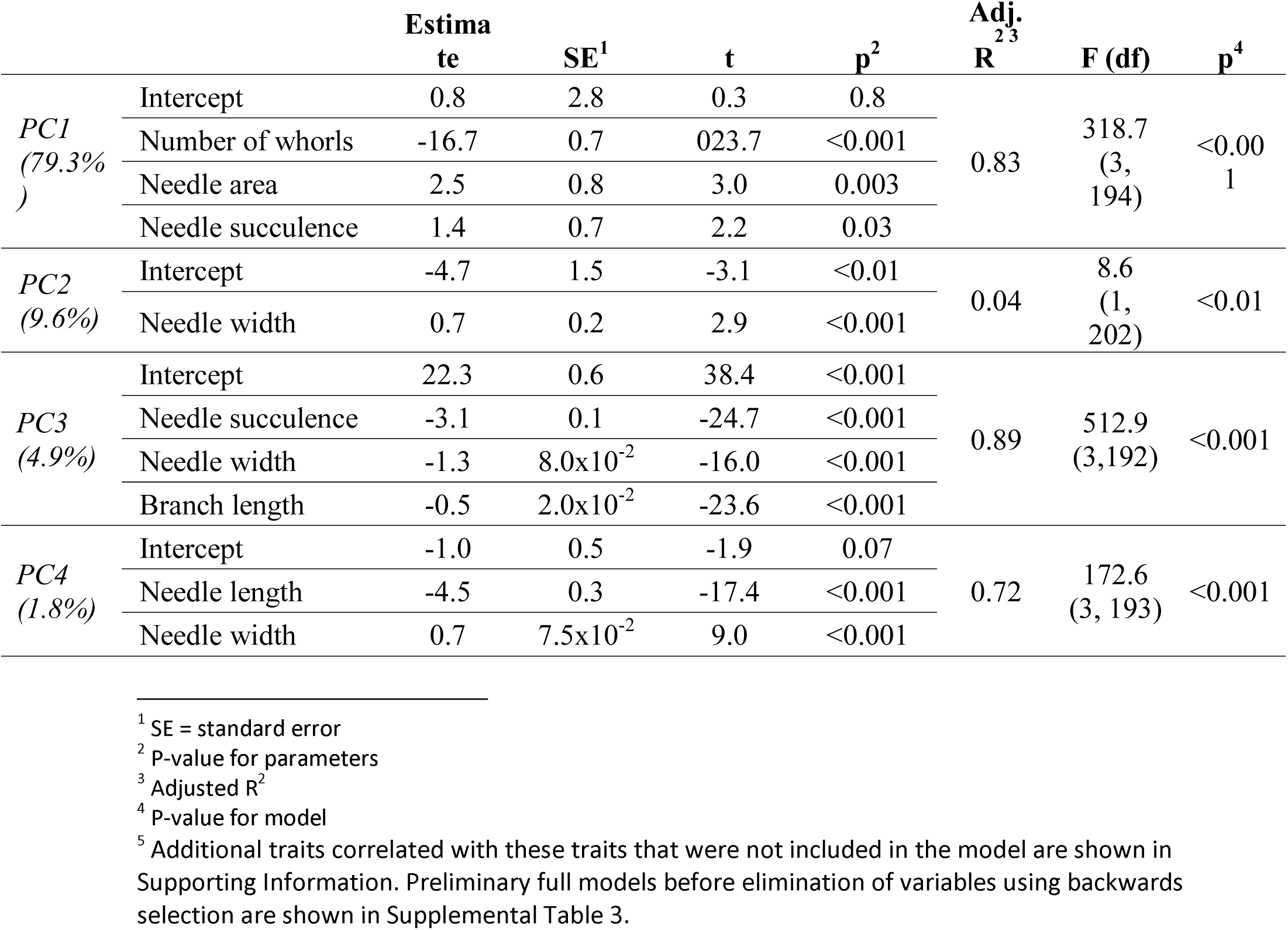
Multiple regression models for traits that explain variation^5^ in the first four PC axes.

For PC2, needle width best explained overall trait variation (Adjusted R^2^=0.04, p=0.004; Table 4, Supporting Information), although it explained relatively little variation. No other traits that were not collinear with needle width were correlated with this axis (where |r|>0.20). Needle area covaried with needle width and showed similar patterns across regions and populations as needle width (Figure 2D). PC3 (which explained 4.9% of overall trait variation) was best explained by needle succulence, needle width, and branch length (Adjusted R^2^=0.88, p<0.001; Table 4, Supporting Information) after removal of survival percentage by backwards selection (ΔAIC=0.0; Supporting Information). PC4 (which explained 1.8% of overall trait variation) was best explained by needle mass per area, needle length, and needle width (Adjusted R^2^=0.72, p<0.001; Table 4, Supporting Information). No traits were removed from the full model.

The first two PC axes primarily differentiated Andes and coastal populations (regions) in their traits (Figure 2B). On average, compared to Andes populations, coastal populations tended to have more whorls (Figure 2C, Supporting Information) and branches (nearly twice as many) as well as branches that are on average 1.5x as long. Number of branches and branch length show similar patterns among regions and populations as number of whorls (shown in Figure 2C). Significant variation is shown within the Andes region among populations as well as within populations in these traits. Additionally, coastal populations tended to have smaller and less succulent needles, with needle area and needle succulence showing similar patterns among regions and populations as needle width (Figure 2D). Needle trait effect sizes were smaller compared to branch architectural traits (Supporting Information).

### Temperature annual range best explained overall region and population trait differences (Q3)

Overall trait differences between regions and among populations were best explained by temperature annual range (TAR), the difference between maximum temperature in the warmest month and minimum temperature in the coldest month (Table 5). Additionally, mean vapor pressure deficit, soil organic carbon, and cation exchange capacity explained small amounts of variation in minor PC axes (Table 5). However, much trait variation remained unexplained by environmental variables.

**Table 5.**
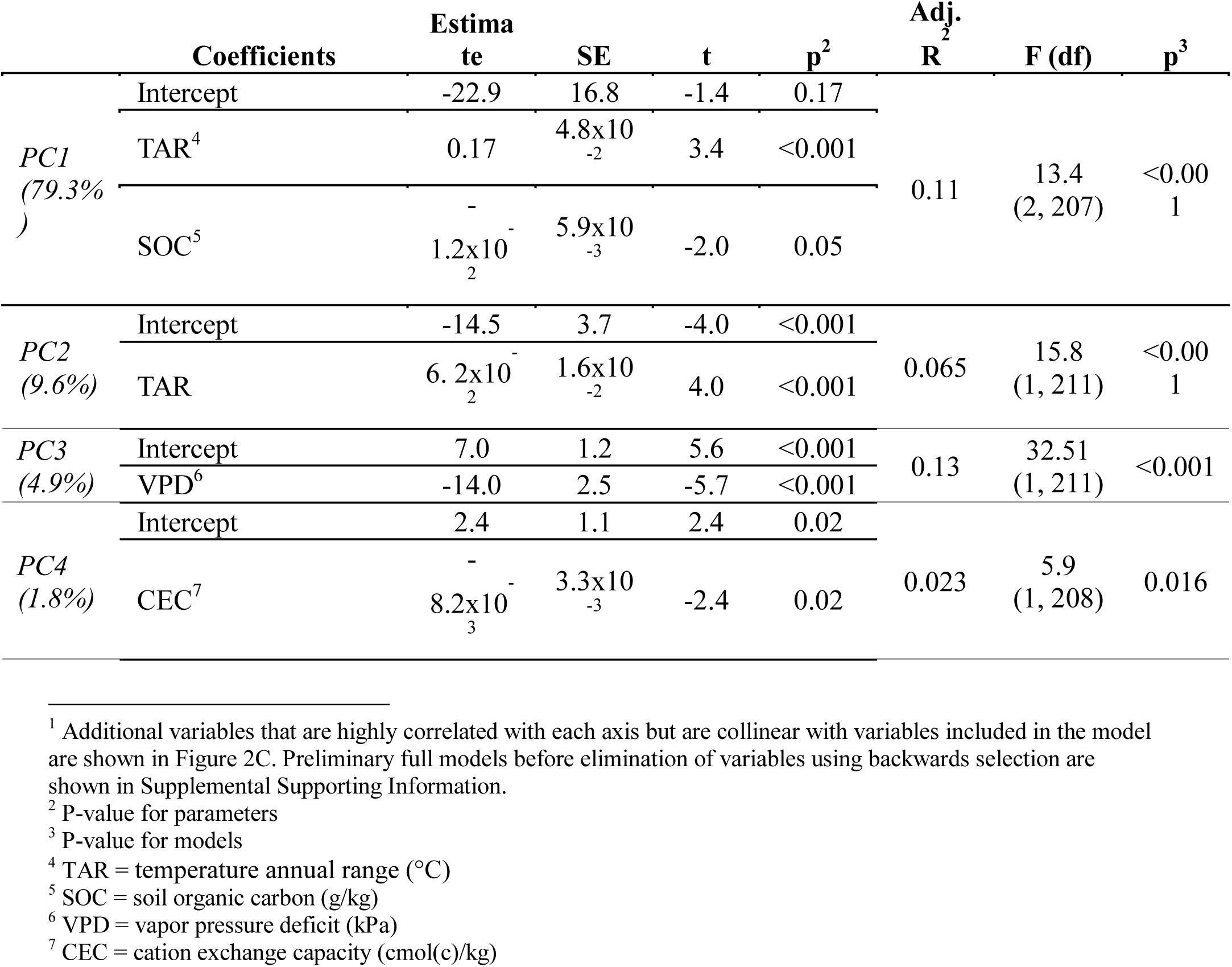
Multiple regression models for environmental variables that explain variation^1^ in the first four PC axes.

For PC1, TAR and SOC together explained 10.6% of variation in PC1 scores (p<0.001, Table 5). Our initial model included SWE, OCD, and Silt, but these variables did not provide additional explanatory power beyond TAR and SOC (ΔAIC=0.0; Supporting Information). Variation in PC2 was also best explained by TAR (although only 6.5% of overall variation was explained; Adjusted R^2^, p<.001, Table 5). For PC2, our initial model included MDR (mean diurnal range) instead of TAR (as Spearman’s r was slightly higher; Supporting Information), but it explained marginally more variation, so we ultimately used TAR for consistency with our model for PC1 (ΔAIC=-0.5; Supporting Information). For PC3, 13% of variation was explained by mean vapor pressure deficit (p<0.001) and for PC4, 2.3% of variation was explained by CEC (p<0.001). For models for PC3 and PC4, no variables were removed from the full models.

Temperature annual range explains 12.0% of all trait variation (p=0.001) and varies significantly among regions and populations (Figure 3, Supporting Information). An additional suite of climate and soil variables covaried with TAR (|r|>0.60) and were thus not included in the multiple regression models (Supporting Information). Overall, coastal populations tend to have smaller temperature annual ranges than Andes populations (Figure 3). This is a result of both higher temperature minimums (−1.58 ± 2.25 vs. -8.00 ± 0.67 °C, p < 0.01) and lower temperature maximums (19.49 ± 2.12 vs. 23.38 ± 1.10 °C, p < 0.001).

**Figure 3.**
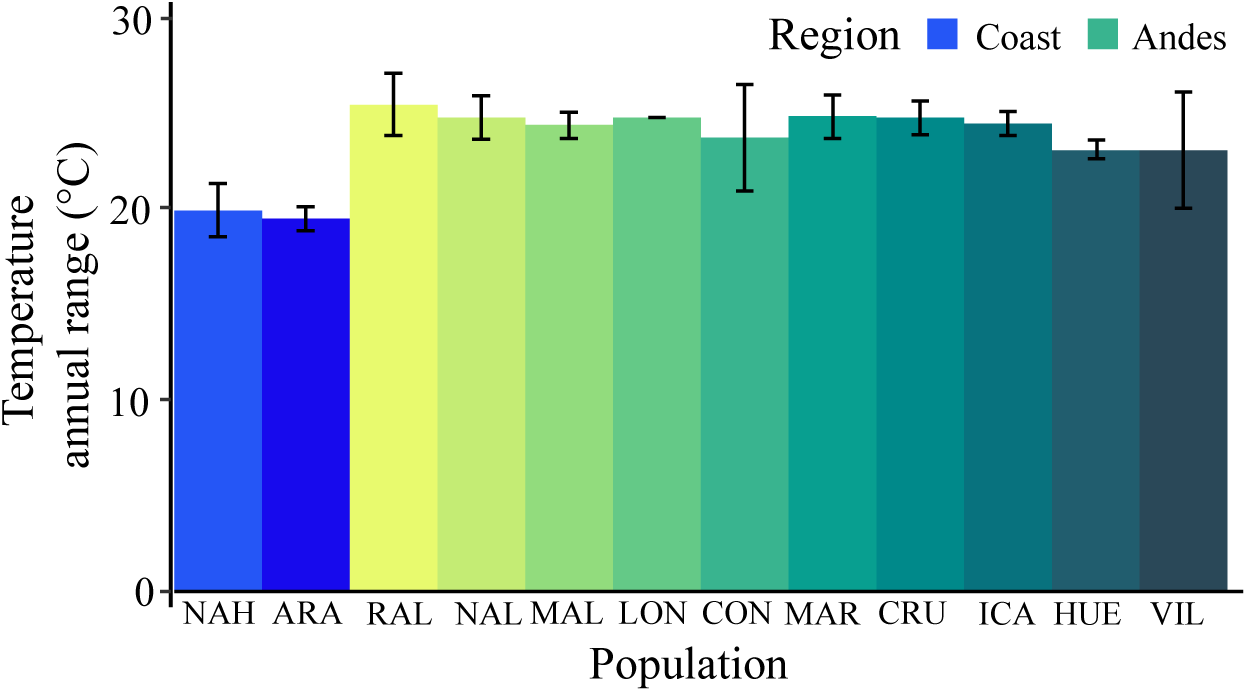
Temperature annual range (TAR, °C) varies significantly among regions and populations. Bars show standard errors. Color categories correspond to region (green = Andes, blue = coast), with gradients by latitude from north (light) to south (dark). Population codes are printed (see Table 1 for more information on populations).

## Discussion

To identify ecotypes for effective restoration and conservation prioritization of threatened species, we must understand patterns of genetic variation in phenotypes across a species’ range, especially in relation to climate and soil variables. Therefore, we asked how populations across the range of *pewen,* an iconic South American conifer species of restoration and conservation concern, varied in a suite of traits between regions and among populations and if this variation was related to climate and soil variables as expected from evidence in other tree species. Our results demonstrate that pewen differs significantly in a suite of traits among and within regions and populations across its range in Chile and that this variation is at least partly explained by climate and soil variables. Temperature annual range, which explained the most trait variation, also explains genomic differentiation in this species (Varas-Myrik et al. 2021). Thus, our results highlight the importance of conserving variation among and within regions, informing conservation strategies and seed sourcing guidelines for restoration.

### Pewen shows differentiation between regions and populations, with high within-population variation

We found clear genetic differentiation in traits between regions. Coastal populations tended to have smaller, less succulent leaves and more branches, while Andes populations tended to have larger, more succulent leaves and fewer branches. Coast to Andes region differences were best explained by temperature annual range, with higher and lower temperature extremes occurring in the Andes region. Thus, we show significant variation in plant traits across the range of pewen in Chile, particularly between the coastal and Andes regions, suggesting that *regional* variation should be conserved. While some trait variation was explained by regional differences, significant variation was also explained by population differences. This suggests that coastal and Andean regions are not only differentiated from each other, but populations within regions are also differentiated from each other and *among-population* variance should be conserved.

Additionally, we showed that a large proportion of variation was unexplained by region or population, suggesting that *within-population* variation should be considered as well.

### Regional differentiation and high within-population variation is consistent with previous assessments of phenotypic and genetic variation in pewen

Our results are consistent with two previous assessments of trait and genetic variation in pewen, which also showed substantial differentiation between coastal and Andes regions and high within-population variation. A phenotypic study of concentrations of alkenes in foliar epicuticular wax, which may contribute to reducing cuticular water loss as an adaptation to drought, revealed differences between coastal and Andes populations (Rafii & Dodd 1998). Although only four populations were used, these authors additionally found high within-population variation in the studied trait. Additional work including nine populations across the coastal and Andes ranges and into pewen’s range in Argentina found that 12% of variation in carbon isotope discrimination and 14% of variation in root:shoot ratio were explained by region (coast, Chilean Andes, Argentinian Andes; Bekessy et al. 2002). These patterns were also corroborated by a study of neutral genetic variation (rather than quantitative genetic variation in traits as assessed here), which found 16% of total variation explained by the region (coast vs.

Chilean Andes; Martín et al. 2014). Two studies using fewer genetic markers and older technology did not detect these trends (Bekessy et al. 2003; Ruiz et al. 2007). Consistent with other work on this species, we found strong evidence of differentiation among mountain ranges (regions). Regional differences in traits could be attributed to genetic isolation; Martín et al. (2014) attributed regional differentiation to geographic isolation among the ranges.

The coastal range is thought to have originated long before the Andes range, and pewen is found on the western slope of the coastal range, a possible barrier to gene flow (as genetic material would have to travel over the coastal range to reach the Andes or vice versa). We also found significant variation that was unexplained by region or population (78% and 89% for PC1 and PC2, respectively; 69% to 95% depending on the trait). In other studies, unexplained trait variation is commonly assumed to be variation maintained within populations (see *Limitations* for further discussion). Within-population variation may be highly important given within-population variation in drought response and subsequent mortality seen in pewen (Puchi et al. 2021).

### High within-population variation and large-scale regional differentiation are common in forest trees

High within-population variation, maintained by gene flow (particularly in wind-pollinated species), is not uncommon for forest trees (Kremer et al. 2012; Alberto et al. 2013). For example, for a small section of the ranges of wind-pollinated ponderosa pine (*Pinus ponderosa)* and Douglas-fir (*Pseudotsuga menziesii)* in Oregon, United States across two mountain ranges with about twice the latitudinal gradient and the same longitudinal gradient as our study, *P. menziesii* but not *P. ponderosa* was differentiated between regions. However, both species showed significantly higher within-population variance compared to among-population variance (Sorensen & Weber 1994). In addition, similar patterns of low among-population variation and high within-population variation in a suite of morphological, phenological, and physiological traits was found in two Northern hemisphere spruces (*Picea glauca, P. engelmannii*) and lodgepole pine (*Pinus contorta)* (Liepe et al. 2016). We did not find additional studies assessing population differentiation in comparable plant traits for other conifers in South America, so we could not compare our results to other local species.

### Temperature annual range best explains overall trait variation

Although temperature annual range explained a limited amount of overall trait variation (12%), this is a substantial amount of variation for just a single climate variable. These findings are consistent with one other study on this species that addressed relationships with environmental variables, where TAR best explained genomic differentiation (Varas-Myrik et al. 2021). Usually, multiple environmental variables play a significant role in explaining multivariate trait variation across populations (Gibson et al. 2019). In our study, TAR primarily explained regional (coast vs. Andes) differences; the magnitude of these differences resulted from both increased minimum and decreased maximum temperatures in the coastal populations, although climate variables associated with temperature maximums tended to be more highly correlated with both PC axes. This suggests that temperature minimums and maximums are both important in shaping population variation in this species. Temperature minimums could explain regional genetic differences in branch traits, as Andes populations experiencing significant frosts (particularly those in the northern part of the range) may not be able to support many large branches due to loss by frost. Temperature maximums could explain regional differences in leaf succulence, with Andes populations experiencing more severe drought having increased succulence (and leaf size) to store water under drought conditions.

Given previous observation of population differentiation in carbon isotope discrimination and cuticular wax alkenes, two traits related to adaptation to arid environments, it is a little surprising that our populations were not differentiated by precipitation or water availability variables. However, meta-analysis shows that most plant traits unrelated to water transport are generally unrelated to precipitation (Griffin-Nolan et al. 2018). Additionally, the lack of explanatory power of water availability variables could be explained by the relatively small precipitation range of this species. Further, additional studies in this species show that differential drought mortality may occur to a greater degree within versus among populations (Puchi et al. 2021).

### Limitations

Our study has four potential limitations. First, selection of adult trees from which to collect seed was not randomized, as seed collection was limited due to availability of seed and ease of access. However, our minimum distance between trees used for seed collection (150m) was greater than that of other studies (50 and 100 m; Rafii and Dodd 1998, Bekessy et al. 2002) and our collection sites within populations varied considerably with respect to topography and microclimate . Thus, we do not feel that this limitation biased results. Second, seedlings were not randomized in the greenhouse, as they were grown in a commercial nursery and subject to procedures therein. However, we anecdotally note that we did not observe any greenhouse effects. Third, we did not replicate within families (trees) in our populations and, therefore, cannot differentiate between within-population variation and error (although it is a common practice in the literature to attribute variance unexplained by population to within-population variation; see Alberto et al. 2013a). If possible, future studies should further replicate within families to account for within-population variation. Finally, there is no information on which plant traits might be adaptive for this species, so we selected traits that have been observed to be important for other species. Thus, we cannot conclude that the variation we identified is adaptive. Future studies are needed to disentangle the traits that are in fact adaptive for this species. Additional work may also consider assessing response to light availability and other environmental factors (which could vary among populations as a result of differences in plant communities).

### Implications for restoration and conservation of pewen in Chile

As ecological restoration commitments ramp up in Chile and beyond, developing science-based resources to guide selection of plant materials is key to maximizing outcomes (Lesica & Allendorf 1999; McKay et al. 2005). In Chile, lack of genetically appropriate seed supply for restoration is a barrier to achieving restoration goals (León-Lobos et al. 2020), although efforts to strengthen seed systems are ongoing (Atkinson et al. 2021). Here, we provide valuable information to complement information on patterns of genomic differentiation (Varas-Myrik et al. 2021)(unpublished data, Ipinza et al. 2021) and assisted migration (Ipinza & Müller-Using 2021) being developed by colleagues to guide conservation prioritization for this species. Given that our data show patterns of variation among as well as within regions and populations, we recommend that restoration efforts aim to collect seed widely within populations across both coastal and Andes mountain ranges, collecting from as many trees within a population as possible to sample within-population diversity (to preserve genetic variation; Kramer and Havens 2009). Additionally, as other studies have concluded, we suggest that managers separate seeds by provenance, particularly avoiding mixing of coastal and Andes seed sources (to avoid maladaptation of seed sources to outplanting sites; Lesica and Allendorf 1999, Broadhurst et al. 2008). We emphasize that conservation of existing and future genetic variation (by widespread seed collection) is necessary to maximize adaptive potential under changing climate, as research indicates this species is at risk within parts of its range (Ipinza & Müller-Using 2021; Varas-Myrik et al. 2021).

Finally, our work sets the stage for the development of seed transfer zones, maps that identify putatively locally adapted ecotypes to guide seed sourcing for restoration (McKay et al. 2005). These resources are needed as provisional zones (which are not species-specific) are generally not sufficient (Gibson & Nelson 2017) and will directly build capacity for restoration in Chile, where collaborators in Chilean management agencies will immediately put them to use. This work, along with additional studies currently in progress by Chilean collaborators, will improve conservation and restoration outcomes for this living fossil species.

## Supporting Information

ANOVA models for all traits with region and population as nested factors (Appendix S1), PC score-trait and -environment correlations (Appendix S2), a figure showing PC score-trait correlations (Appendix S3), full multiple regression models for traits (Appendix S4), means and SEs for traits differentiated by region (Appendix S5), full multiple regression models for environmental variables (Appendix S6), ANOVA models for temperature annual range with region and population as nested factors (Appendix S7), and a figure showing PC score-environmental variable correlations (Appendix S7) are available online. The authors are solely responsible for the content and functionality of these materials. Queries (other than absence of the material) should be directed to the corresponding author.

## Impact Statement

Differentiation in key traits within and among regions guides restoration and conservation efforts for the iconic Chilean tree *Araucaria araucana*.

## Acknowledgements

This study was designed by MM and CN. JG and IR selected seed collection sites and facilitated seedling germination and growth with a commercial greenhouse. MM, PD, OT, N. Gutierrez, and I. Arguto collected data. MM analyzed the data and wrote the manuscript. All authors helped with review of the manuscript. Additionally, we acknowledge the Sistema de Monitoreo de Ecosistemas Forestales Nativos and the Instituto Forestal in Chile for financial and logistical support of this project. We are grateful to our Mapuche Pewenche partners who contributed to seed collection for many of our populations. Finally, we thank CMPC and the Carlos Douglass Nursery for assistance in germinating and growing seedlings. We greatly appreciate the support of Dr. Patricia Saez Delgado and her lab for providing lab space for data collection. MM was supported by NSF DGE-184053 and additional scholarships from the University of Montana Franke College of Forestry and Conservation.

