## Supporting Information for "Trait variation between and within Andes and coastal mountain ranges in the iconic South American tree *Araucaria araucana* in Chile"

**Appendix 1.** ANOVA tables for traits that varied significantly among regions or populations and were included in PCA (top) and traits that did not vary significantly among regions or populations and were not included in PCA (bottom).

| **Traits included in Multiple Regression Models** | | | | | | | | | |
| --- | --- | --- | --- | --- | --- | --- | --- | --- | --- |
|  |  | **df** | **SS** | **MSE** | **F** | **p** | **F (df)** | **Adj. R^2^** | **p** |
| *Needle width* | Region | 1 | 26.7 | 26.7 | 34.5 | <0.001 | 9.3 (11,192) | 0.31 | <0.001 |
|  | Population | 10 | 52.7 | 5.3 | 6.8 | <0.001 |  |  |  |
|  | Residuals | 192 | 148.5 | 0.8 |  |  |  |  |  |
| *Needle area* | Region | 1 | 15.6 | 15.6 | 69.5 | <0.001 | 7.6 (11, 194) | 0.26 | <0.001 |
|  | Population | 10 | 3.2 | 3.2 | 1.4 | 0.2 |  |  |  |
|  | Residuals | 194 | 43.6 | 0.2 |  |  |  |  |  |
| *Needle succulence* | Region | 1 | 11.0 | 11.0 | 28.1 | <0.001 | 4.6 (11, 194) | 0.16 | <0.001 |
|  | Population | 10 | 8.6 | 0.9 | 2.2 | 0.02 |  |  |  |
|  | Residuals | 194 | 76.0 | 0.4 |  |  |  |  |  |
| *Number of whorls* | Region | 1 | 9.5 | 9.5 | 27.5 | <0.001 | 4.6 (11, 193) | 0.16 | <0.001 |
|  | Population | 10 | 8.0 | 0.8 | 2.3 | 0.01 |  |  |  |
|  | Residuals | 193 | 66.9 | 0.3 |  |  |  |  |  |
| *Needle thickness* | Region | 1 | 0.5 | 0.5 | 26.3 | <0.001 | 5.3 (11, 194) | 0.19 | <0.001 |
|  | Population | 10 | 0.6 | 0.1 | 3.2 | 0.001 |  |  |  |
|  | Residuals | 194 | 3.7 | 1.9x10^-2^ |  |  |  |  |  |
| *Needle length* | Region | 1 | 14.9 | 14.9 | 35.3 | <0.001 | 3.9 (11, 192) | 0.13 | <0.001 |
|  | Population | 10 | 3.1 | 0.3 | 0.7 | 0.7 |  |  |  |
|  | Residuals | 192 | 81.2 | 0.4 |  |  |  |  |  |
| *Number of branches* | Region | 1 | 24.7 | 24.7 | 20.0 | <0.001 | 3.8 (11, 193) | 0.13 | <0.001 |
|  | Population | 10 | 26.7 | 2.7 | 2.2 | 0.02 |  |  |  |
|  | Residuals | 193 | 237.5 | 1.23 |  |  |  |  |  |
| *Needle mass per area* | Region | 1 | 0.6 | 0.6 | 8.0 | <0.01 | 3.2 (11, 194) | 0.10 | <0.001 |
|  | Population | 10 | 2.1 | 0.21 | 2.7 | <0.01 |  |  |  |
|  | Residuals | 194 | 15.3 | 7.9x10^-2^ |  |  |  |  |  |
| *Branch length* | Region | 1 | 133.2 | 133.2 | 11.3 | <0.001 | 2.7 (11, 192) | 0.09 | 0.002 |
|  | Population | 10 | 222.4 | 22.2 | 1.9 | 0.05 |  |  |  |
|  | Residuals | 192 | 2268.7 | 11.8 |  |  |  |  |  |
| *Percent survival* | Region | 1 | 0.2 | 0.2 | 14.0 | <0.001 | 2.2 (11, 201) | 0.06 | 0.014 |
|  | Population | 10 | 0.1 | 1.5x10^-2^ | 1.1 | 0.40 |  |  |  |
|  | Residuals | 201 | 2.8 | 0.01 |  |  |  |  |  |
| *Percent germination* | Region | 1 | 2.1x10^-2^ | 2.1x10^-2^ | 1.7 | 0.2 | 2.0 (11, 201) | 0.05 | 0.03 |
|  | Population | 10 | 0.3 | 2.510^-2^ | 2.0 | 0.03 |  |  |  |
|  | Residuals | 192 | 2.4 | 1.2x10^-2^ |  |  |  |  |  |
| **Traits not Included in Multiple Regressions** | | | | | | | | | |
|  |  | **df** | **SS** | **MSE** | **F** | **p** |  |  | **df** |
| *Aboveground biomass* | Region | 1 | 1.4 | 1.4 | 0.5 | 0.5 | *Aboveground biomass* | Region | 1 |
|  | Population | 10 | 11.5 | 1.2 | 0.5 | 0.9 |  | Population | 10 |
|  | Residuals | 178 | 457.4 | 2.6 |  |  |  | Residuals | 178 |
| *Belowground biomass* | Region | 1 | 1.0x10^-2^ | 7.5 x10^-3^ | 2.0 x10^-2^ | 0.9 | *Belowground biomass* | Region | 1 |
|  | Population | 10 | 1.81 | 0.2 | 0.6 | 0.8 |  | Population | 10 |
|  | Residuals | 178 | 56.3 | 0.3 |  |  |  | Residuals | 178 |
| *Stem length* | Region | 1 | 3.4 | 3.4 | 0.4 | 0.5 | *Stem length* | Region | 1 |
|  | Population | 10 | 50.7 | 5.1 | 0.5 | 0.9 |  | Population | 10 |
|  | Residuals | 189 | 1783.3 | 9.4 |  |  |  | Residuals | 189 |
| *Basal diameter* | Region | 1 | 1.8 | 1.8 | 1.5 | 0.2 | *Basal diameter* | Region | 1 |
|  | Population | 10 | 16.2 | 1.6 | 1.4 | 0.2 |  | Population | 10 |
|  | Residuals | 186 | 215.0 | 1.2 |  |  |  | Residuals | 186 |

**Appendix 2.** Correlations between PC scores and traits or environmental variables. Provided online as a separate Excel file.

**Appendix 3.** A suite of traits are correlated with PC axis scores, contributing to explaining the largest proportion of overall trait variation among populations across the range of pewen in Chile. Spearman’s r correlations among A) PC1 axis scores, B) PC2 axis scores, C) PC3 axis scores, D) PC4 axis scores and traits, where |r|>0.20. Large dashed gray line denotes r=0 and small dashed grey lines are where |r|=0.50. Symbols denote p-values for Spearman’s correlations (see legend). Trait units are shown in Table 2.

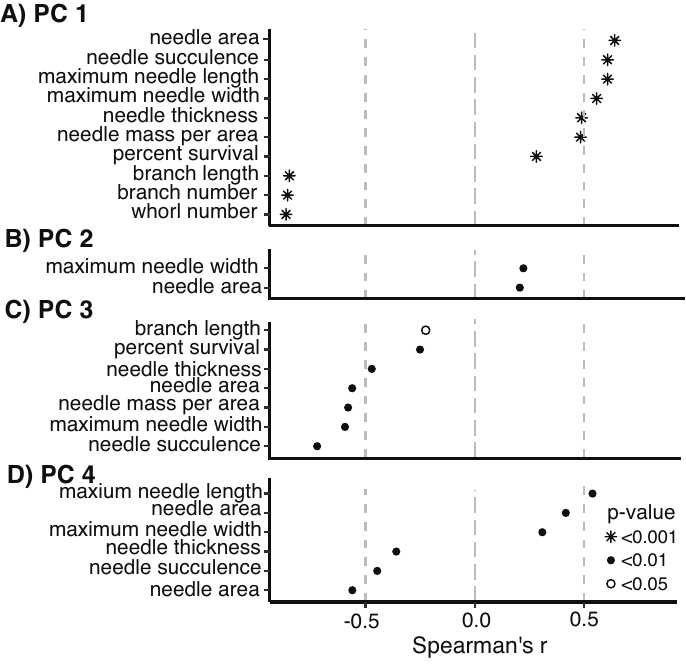

**Appendix 4.** ANOVA outputs from full PC score-trait models before nonsignificant traits were excluded. Reduced models which were used in this manuscript and associated ΔAIC values are included in Table 4. For PC2 and PC4, full models are used (Table 4). Traits which were included in final models are italicized.

|  | **Coefficients** | **Estimate** | **SE** | **t** | **p** | **Adj. R^2^** | **F (df)** | **p** | **AIC (Δ AIC)** |
| --- | --- | --- | --- | --- | --- | --- | --- | --- | --- |
| *PC1*  *(79.3%)* | Intercept | 2.0 | 3.6 | 0.6 | 0.6 | 0.83 | 238  (4, 193) | <0.001 | 1237.1 (1.7) |
|  | *Number of whorls* | *-16.7* | *0.7* | *-23.6* | *<0.001* |  |  |  |  |
|  | *Needle area* | *2.6* | *0.8* | *3.0* | *0.003* |  |  |  |  |
|  | *Needle succulence* | *1.5* | *0.7* | *2.2* | *0.03* |  |  |  |  |
|  | Percent survival | -1.7 | 3.3 | -0.5 | 0.6 |  |  |  |  |
| *PC3*  *(4.9%)* | Intercept | 22.8 | 0.7 | 32.6 | <0.001 | 0.89 | 387  (4, 191) | <0.001 | 572.6  (0.0) |
|  | Needle succulence | -3.1 | 0.1 | -24.2 | <0.001 |  |  |  |  |
|  | Needle width | -1.3 | 0.1 | -15.8 | <0.001 |  |  |  |  |
|  | Branch length | -0.5 | 2.2x10^-2^ | -23.7 | <0.001 |  |  |  |  |
|  | Percent survival | -0.9 | 0.6 | -1.4 | 0.2 |  |  |  |  |

**Appendix 5.** Means ± standard errors for key branch and needle traits that are differentiated regionally.

| **Trait** | **Coastal populations** | **Andes populations** |
| --- | --- | --- |
| *Number of whorls* | 1.1 ± 0.11 | 0.6 ± 0.05 |
| *Number of branches* | 1.9 ± 0.23 | 0.98 ± 0.08 |
| *Branch length (cm)* | 5.3 ± 0.23 | 3.3 ± 0.08 |
| *Needle width (mm)* | 5.4 ± 0.12 | 6.3 ± 0.08 |
| *Needle succulence (mg/cm^2^)* | 3.50 ± 0.07 | 4.1 ± 0.05 |

**Appendix 6.** ANOVA outputs from full PC score-environmental variable models before nonsignificant traits were excluded. Reduced models which were used in this manuscript and associated ΔAIC values are included in Table 4. For PC3 and PC4, full models are used (Table 4). Traits which were included in final models are italicized.

|  | **Coefficients** | **Estimate** | **SE** | **t** | **p** | **Adj. R^2^** | **F (df)** | **p** | **AIC (Δ AIC)** |
| --- | --- | --- | --- | --- | --- | --- | --- | --- | --- |
| *PC1*  *(79.3%)* | Intercept | 30.1 | 45.9 | 0.7 | 0.5 | 0.09 | 4.3  (7, 202) | <0.001 | 1653.9 (0.0) |
|  | TAR | -4.6x10^-4^ | 0.1 | -3.6x10^-1^ | 1.0 |  |  |  |  |
|  | SWE | -3.3x10^-2^ | 7.3x10^-2^ | -0.4 | 0.7 |  |  |  |  |
|  | Silt | 3.5x10-^2^ | 2.4x10^-2^ | 1.5 | 0.1 |  |  |  |  |
|  | SOC | -1.2x10^-2^ | 6.2x10^-3^ | -1.9 | 0.05 |  |  |  |  |
|  | Clay | ­-1.7x10^-2^ | 2.8x10^-2^ | -0.6 | 0.5 |  |  |  |  |
|  | VAP | -24.93 | 23.7 | -1.1 | 0.3 |  |  |  |  |
|  | OCD | -5.9x10^-3^ | 2.6x10-2 | -0.2 | 0.8 |  |  |  |  |
| *PC2*  *(9.6%)* | Intercept | -14.0 | 3.6 | -3.9 | <0.001 | 0.06 | 15.28  (1, 211) | <0.001 | 1237.2  (-0.52) |
|  | MDR | 0.1 | 2.7x10-^2^ | 3.9 | <0.001 |  |  |  |  |

**Appendix 7.** ANOVA model outputs for differentiation in temperature annual range (TAR, °C) by region and population (nested within region).

|  |  | **df** | **SS** | **MSE** | **F** | **p** | **Adj. R^2^** | **p** |
| --- | --- | --- | --- | --- | --- | --- | --- | --- |
| *TAR* | Region | 1 | 66900 | 66900 | 28271 | <0.001 | 0.99 | <0.001 |
|  | Population | 10 | 11003 | 1100 | 465 | <0.001 |  |  |
|  | Residual | 201 | 476 | 2 |  |  |  |  |

**Appendix 8.** A suite of environmental variables are correlated with PC axis scores, explaining the largest proportion of overall trait variation among regions and populations across the range of pewen in Chile. **A)** Spearman’s r correlations among PC1 axis scores and environmental variables where |r|>0.20. **B)** Spearman’s r correlations among PC2 axis scores and environmental variables, where |r|>0.20. Large dashed gray line denotes r=0 and small dashed grey lines are where |r|=0.50. Symbols denote p-values for Spearman’s correlations (see legend). Full environmental variable names and units are included in Table 3).

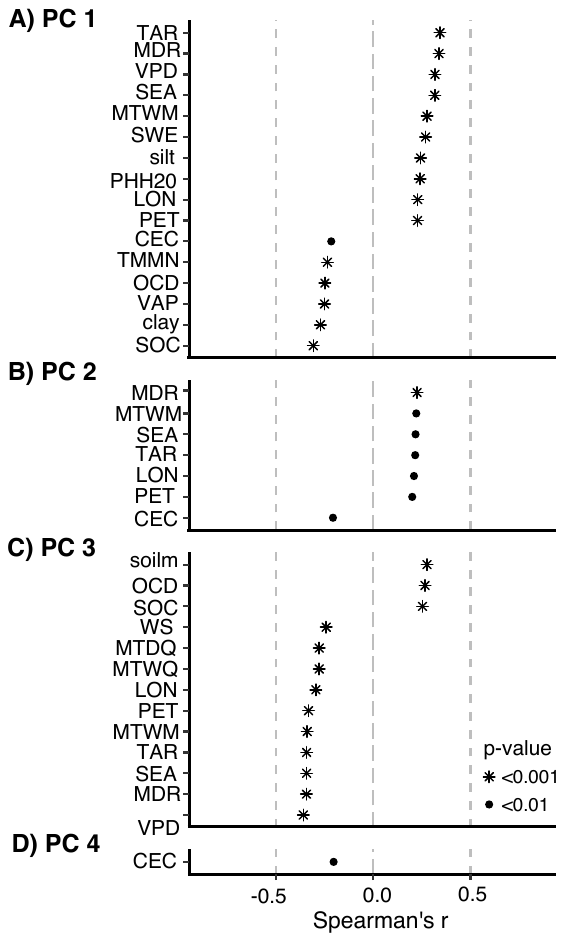
